# The effectiveness of acellular nerve allografts compared to autografts in animal models: a systematic review and meta-analysis

**DOI:** 10.1101/2022.12.06.519361

**Authors:** Berend O. Broeren, Caroline A. Hundepool, Ali H. Kumas, Liron S. Duraku, Erik T. Walbeehm, Carlijn R. Hooijmans, Dominic M. Power, J. Michiel Zuidam, Tim De Jong

## Abstract

**Background:** Treatment of nerve injuries proves to be a worldwide clinical challenge. Acellular nerve allografts are suggested to be a promising alternative for bridging a nerve gap to the current gold standard, an autologous nerve graft.

**Objective:** To systematically review the efficacy of the acellular nerve allograft, its difference from the gold standard (the nerve autograft) and to discuss its possible indications.

**Material and methods:** PubMed, Embase and Web of Science were systematically searched until the 4th of January 2022. Original peer reviewed paper that presented 1) distinctive data; 2) a clear comparison between not immunologically processed acellular allografts and autologous nerve transfers; 3) was performed in laboratory animals of all species and sex. Meta analyses and subgroup analyses (for graft length and species) were conducted for muscle weight, sciatic function index, ankle angle, nerve conduction velocity, axon count diameter, tetanic contraction and amplitude using a Random effects model. Subgroup analyses were conducted on graft length and species.

**Results:** Fifty articles were included in this review and all were included in the meta-analyses. An acellular allograft resulted in a significantly lower muscle weight, sciatic function index, ankle angle, nerve conduction velocity, axon count and smaller diameter, tetanic contraction compared to an autologous nerve graft. No difference was found in amplitude between acellular allografts and autologous nerve transfers. Post hoc subgroup analyses of graft length showed a significant reduced muscle weight in long grafts versus small and medium length grafts. All included studies showed a large variance in methodological design.

**Conclusion:** Our review shows that the included studies, investigating the use of acellular allografts, showed a large variance in methodological design and are as a consequence difficult to compare. Nevertheless, our results indicate that treating a nerve gap with an allograft results in an inferior nerve recovery compared to an autograft in seven out of eight outcomes assessed in experimental animals. In addition, based on our preliminary post hoc subgroup analyses we suggest that when an allograft is being used an allograft in short and medium (0-1cm, > 1-2cm) nerve gaps is preferred over an allograft in long (> 2cm) nerve gaps.

## Introduction

Peripheral nerve injuries affect 2,8% of all trauma cases, and despite surgical repair, they often result in deterioration of quality of life for these patients. (1, 2) (3) In some injuries, a segmental loss of a peripheral nerve occurs after trauma or tumor excision for example. (4, 5)

The current gold standard for the surgical repair of a peripheral nerve injury that cannot be directly coaptated is a nerve autograft. The sural nerve is the most commonly used because it supplies a consistent source of graft material and is anatomically accessible. (6) However, this procedure has several limitations. The length of donor nerve available and its limited diameter are often insufficient to achieve a complete reconstruction of multiple or significant segmental defects. Besides, the procedure may cause considerable donor site morbidity, such as pain and loss of sensation. (7-9)

Several techniques have been investigated to replace the nerve autograft, including allografts, biological conduits and synthetic conduits. (10-12) All with their benefits and drawbacks. Of these options, the acellular allograft seems the most promising. (13) These grafts provide the needed internal structural and molecular composition of the extracellular matrix in a cell-free scaffold, which supports nerve regeneration while retaining a nonimmunogenic nature. (14) This procedure however, has some drawbacks as well, including uncertain histocompatibility and ethical and legal concerns.

A variety of methods have been studied to prepare an acellular allograft such as cold preservation, freeze-thaw cycling, chemical detergent and enzymatic preparations, lyophilization and irradiation. (15-17) There is only one such method that is FDA approved available to surgeons produced by AxoGen, Inc., Alachua, Florida. AxoGen develops its human allografts by combining proprietary detergent processing and gamma irradiation which removes cellular remnants while minimizing the microstructural damage. Next to that, chondroitin sulfate proteoglycan is enzymatic removed from the endoneural tube system to advance axon regeneration. (18) Several additions and alterations to this method have been researched. However, at this moment these techniques are not clinically available (Fig 2. shows an approximate overview of these methods).

There is little clinical and experimental evidence about the difference in outcomes between an acellular allograft and an autograft. Therefore, A systematic review of experimental studies was conducted to investigate the efficacy of the acellular nerve allograft, its difference from the gold standard (the nerve autograft) and to discuss the possible indications for the use of an allograft.

## Material and methods

### Research protocol

Before starting this systematic review, a protocol was defined in advance and registered in an international database (PROSPERO, registration number CRD42020186451).

### Search strategy

PubMed (Medline), Embase (OVID) and Web of Science were systematically searched to identify all original articles. The search contained studies up to the 4th of January 2022. Search terms included ‘nerve reconstruct’, ‘nerve transfer’, ‘nerve graft’, ‘allograft’, ‘allogeneic’, ‘acellular’, ‘decellularize’ and their synonyms in abstract and title fields (see S1 Table for the complete search strategy). To identify all animal studies, the SYRCLE search filters were used. (19, 20) Endnote (Clarivate Analytics, Pennsylvania, USA) was used to remove duplicates. Two authors (BOB and AK) performed the screening process independently using Rayyan web tool. (21) All titles and abstracts were screened to determine their relevance by utilizing the pre-established inclusion and exclusion criteria. Reference lists of the remaining studies were screened manually for potentially relevant new studies. Full text screening of all relevant articles was done by two reviewers for final selection. Divergences were solved by consensus discussion. Any remaining divergences were solved by consulting TDJ as a third reviewer.

### Inclusion and exclusion criteria

We included an original peer reviewed paper 1) that presented distinctive data; 2) made a clear comparison between not immunologically processed acellular allografts and autologous nerve transfers; 3) was performed in laboratory animals of all species and sex; 4) investigated the effect of acellular allografts on motor outcomes: sciatic function index, muscle weight (gram), ankle angle (degrees), electrophysiology (nerve conduction velocity (ms/s), amplitude (mA) or latency (ms)) and sensory outcomes: hot-cold testing, pin-prick testing, Semmes-Weinstein testing and histomorphometry (axon count and diameter). No publication date restriction was applied.

### Critical appraisal

Two authors (BOB and AK) independently assessed the risk of bias using the SYRCLE’s tool for assessing the risk of bias for animal studies. This appraisal was subsequently merged by consensus and disagreements were solved by discussion. (22) A “yes” indicating a low risk of bias, a “no” indicating a high risk of bias or a “?” indicating an unknown risk of bias was scored for all criteria. We determined selective outcome reporting by establishing if all outcome measures stated in the material and methods section were also reported in the results. Baseline characteristics were: species, age and weight. We included two items to overcome the problem of judging to many items as “unclear risk of bias: reporting on any measure of randomization and reporting on any measure of blinding. For these two questions a “yes” indicates reported and a “no” indicates not reported.

### Data extraction

From the included studies, both reviewers (BOB and AK) extracted the data in duplicate. The descriptive data included: first author’s name, the year of publication, studied species, sex, total number of animals, number of grafts, studied nerve, studied muscle, graft size and time points. The mean, standard deviation (SD) and total number of subjects (n) were recorded for all outcomes. In case multiple locations per nerve were reported, we used the most distal segment of the graft. When the SEM was reported, it was recalculated to SD (SD = SEM x √n). If data were only presented graphically, Universal Desktop Ruler software (https://avpsoft.com/products/udruler/) was used by two reviewers independently to measure a fair estimation of the presented data, after that the mean of these two independent measurements was used. We attempted to contact the authors for additional information in case relevant data were missing

### Statistical analysis

Comprehensive Meta-Analysis (CMA version 3.3) was used to analyze all data. The standardized mean difference (SMD) and 95% confidence interval (95% CL) for all outcome measurements comparing acellular allografts and conventional autografts were calculated with Hedges’ g correction. A random effects model was applied, which takes the accuracy of independent studies and the variation among studies into account and weighs all studies accordingly. I^2^ was used to asses heterogeneity. In case a study reported results at different time points using the same experimental group, these results were pooled to obtain an overall SMD with Hedges’g correction using a random-effects model and variance. Subgroup analyses were conducted post hoc for species (rat, rabbit, monkey and dog) and graft lengths (0-1 cm, > 1-2 cm and > 2 cm). We only interpreted the results of subgroup analysis when groups consisted of 5 or more individual studies.

To detect publication bias funnel plots were created and evaluated on symmetry using Egger’s regression and Trim and Fill analysis, if there were at least 15 or more independent studies per outcome. We plotted the SMD against a sample size-based precision estimate(1/√(n)), because SMDs may cause funnel plot distortion.

A sensitivity analysis was performed to assess the robustness of our findings. The impact of excluding studies published before 2008 and studies that used animals as their own control was evaluated.

## Results

### Study selection process

The systematic literature search presented in S1 Table yielded 1191 unique references (Fig 1. shows a consort flow chart). After title abstract screening, 136 studies met the selection criteria. Finally, after studying the full-text articles, 50 studies were included in the review and meta-analyses.

**Fig 1.**
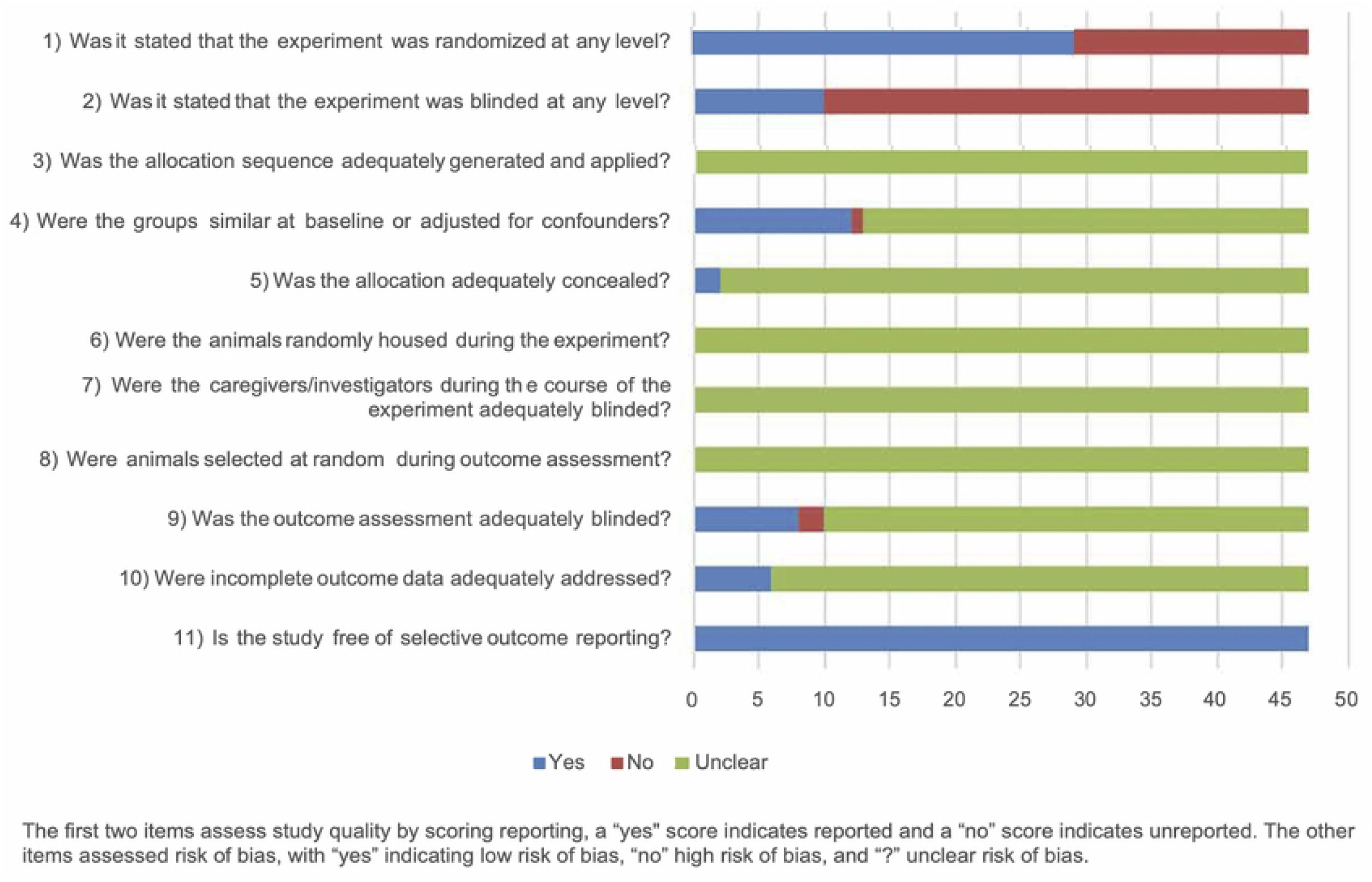
Flow chart of the study selection.

**Fig 2.**
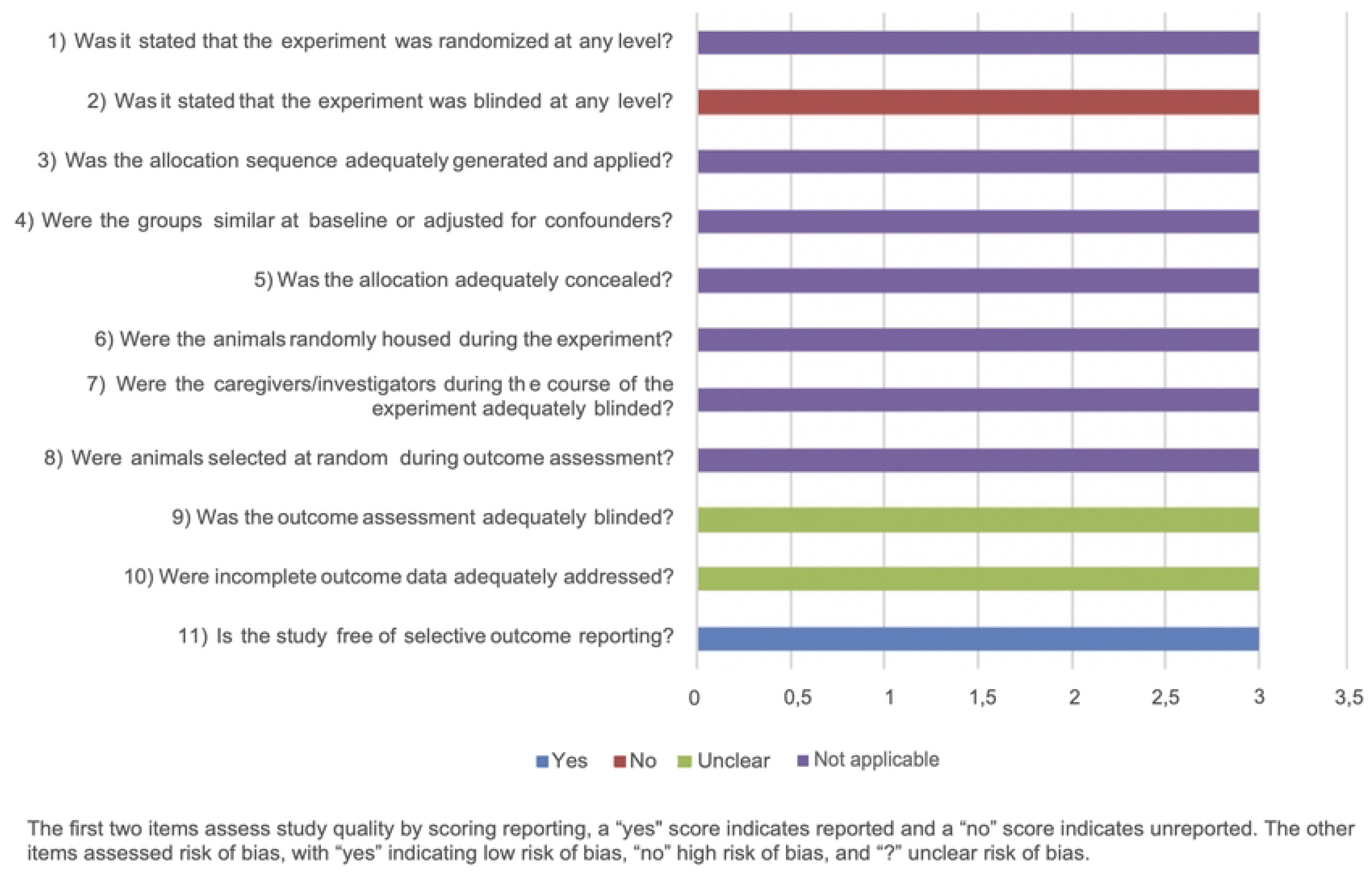
Global overview decellularization methods.

### Study quality and risk of bias

The general results of our risk of bias assessment of the included references are shown in Fig 3. Poor reporting of essential methodological details in most animal experiments resulted in an unclear risk of bias in most studies. In particular reporting about any randomization and blinding measures taken at any level was 64% (32 out of 50 publications). Assessment of the risk of bias was done separately for the 3 studies that used animals as their own control because some aspects were not applicable (Fig 4).

**Fig 3.**
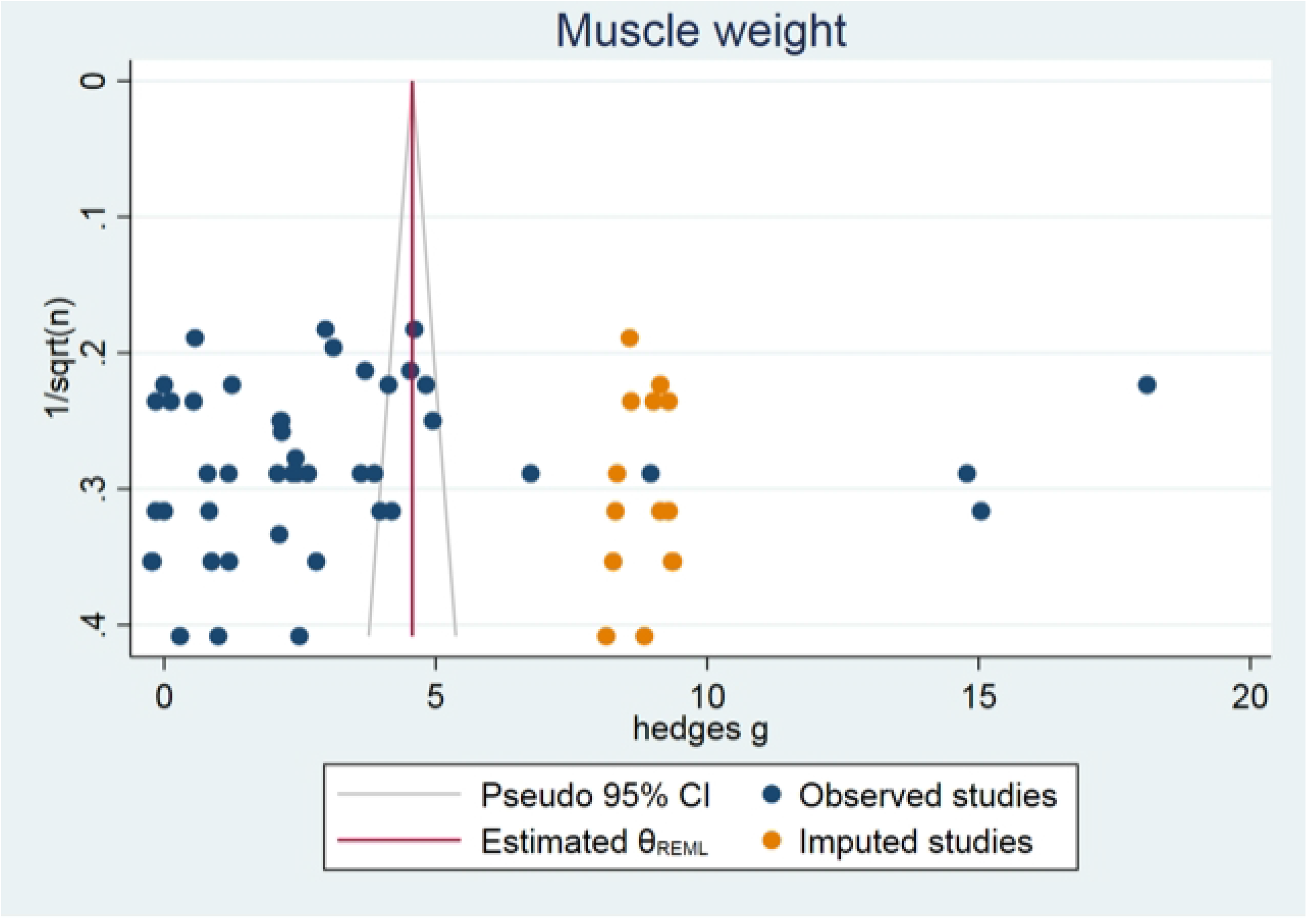
Results of the risk of bias assessment of 47 included studies in this systematic review. The first two items assess study quality by scoring reporting, a “yes” score indicates reported and a “no” score indicates unreported. The other items assessed risk of bias, with “yes” indicating low risk of bias, “no” high risk of bias, and “?” unclear risk of bias.

**Fig 4.**
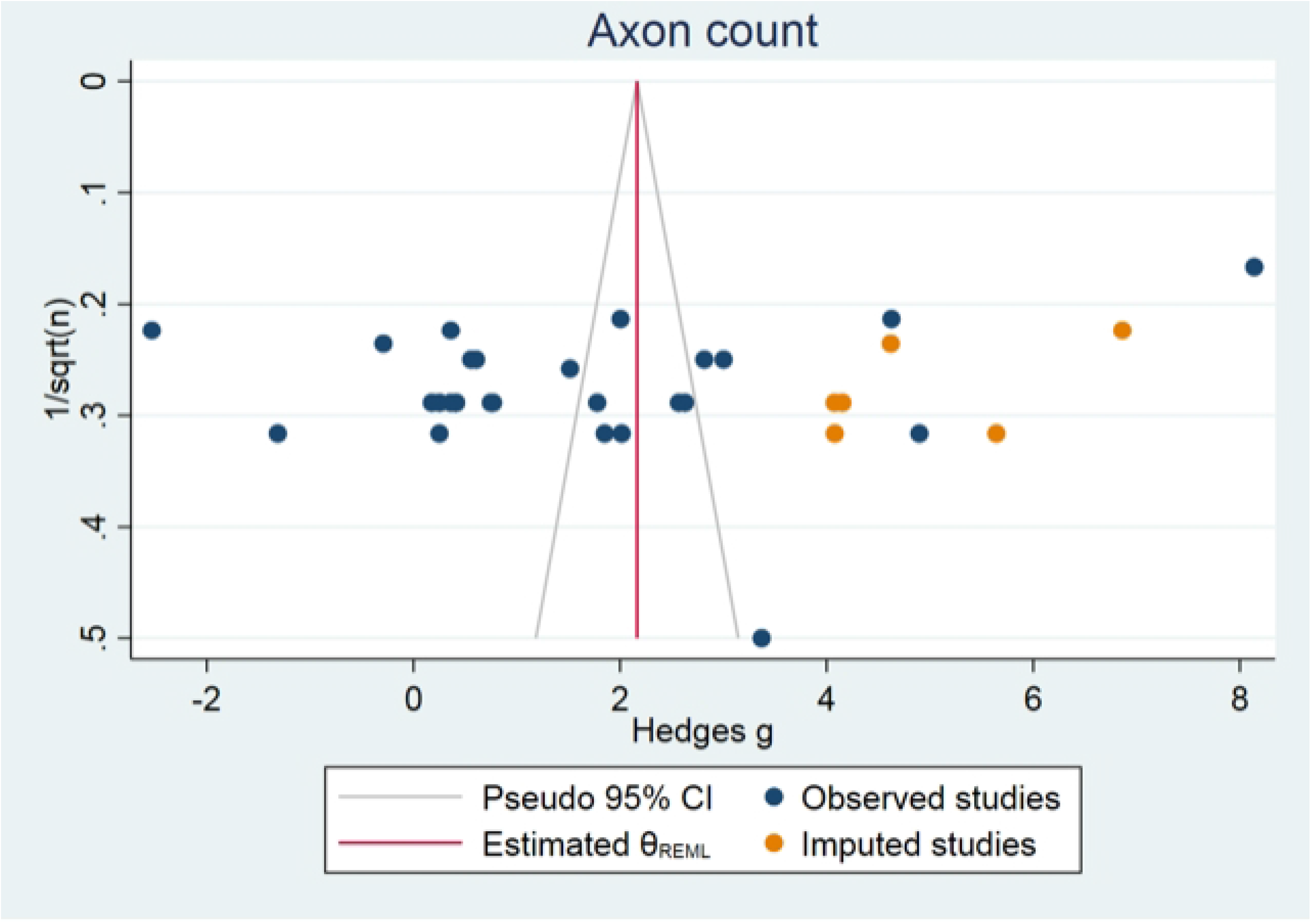
Results of the risk of bias assessment of the 3 included studies in this systematic review where animals were their own control group. The first two items assess study quality by scoring reporting, a “yes” score indicates reported and a “no” score indicates unreported. The other items assessed risk of bias, with “yes” indicating low risk of bias, “no” high risk of bias, and “?” unclear risk of bias.

### Study characteristics

A summary of the characteristics of the 50 included publications is shown in Table 1. (23-71) The characteristics per outcome measurement are depicted in the Appendix. The most commonly used specie was rat (80%), followed by rabbit (6%), monkey (6%), dog (6%) and mice (2%). Gender was not reported in 20% (10 of 50) of the publications. Out of the remaining studies 33 used males, 2 used females and in 5 both sexes were used. Different nerves were used with the sciatic nerve being the most common (80%), followed by the peroneal nerve (6%), facial nerve (4%), radial nerve (4%), ulnar nerve (2%), tibial nerve (2%) and femoral nerve (2%).

**Table 1.**
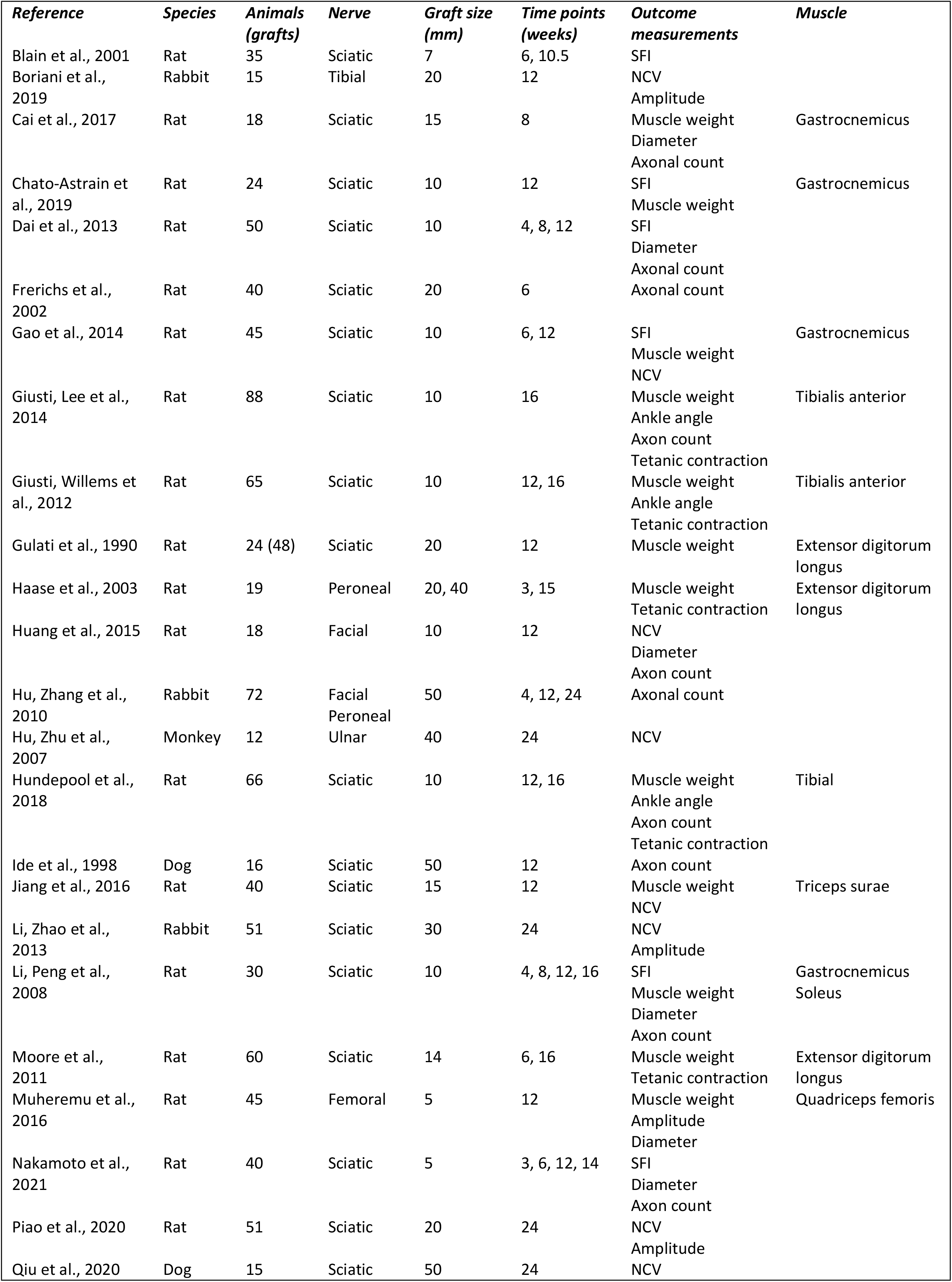
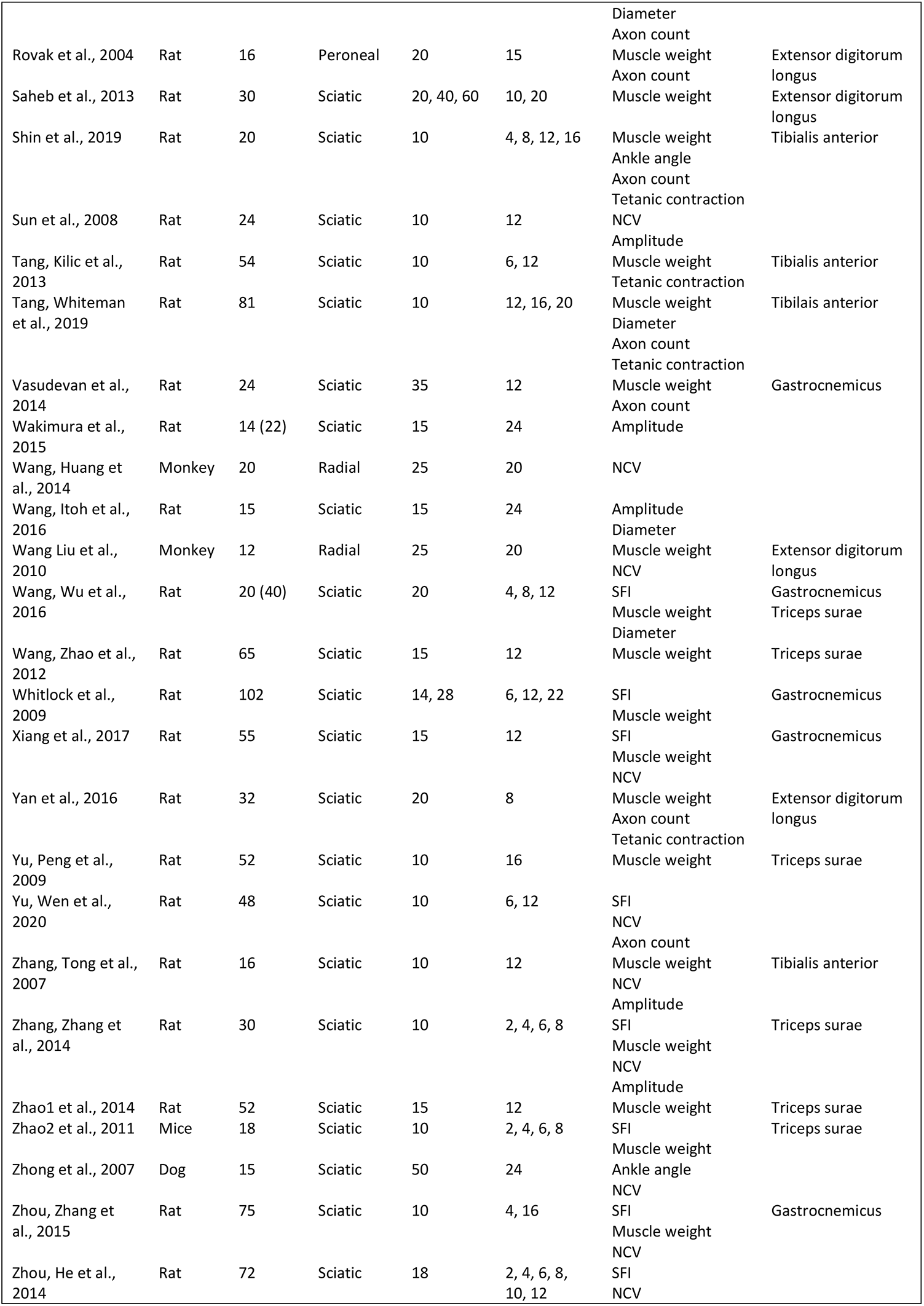
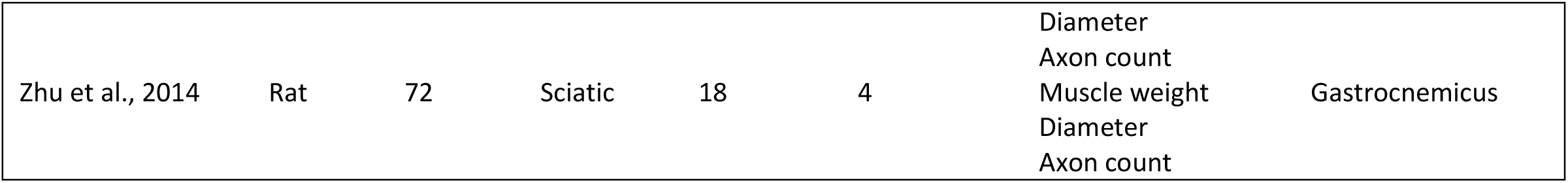
The characteristics of all 50 included references.

### Overall analysis

Overall analysis showed a significant lower muscle weight, sciatic function index, tetanic contraction, nerve conduction velocity and smaller ankle angle, axon count diameter after treatment with an acellular allograft compared to an autograft (Shown in Table 2). No significant difference in amplitude was found.

**Table 2.** Summary of the overall analyses.

| Outcome measurement | SMD (Hedges g) | 95% confidence interval | $I^2$ | No. of comparisons | No. of studies |
| --- | --- | --- | --- | --- | --- |
| Muscle weight | -2.39 | -1.85 to -2.93 | 86% | 45 | 31 |
| Sciatic function index | -1.59 | -0.44 to -2.73 | 93% | 15 | 14 |
| Tetanic contraction | -0.54 | -0.13 to -0.95 | 62% | 14 | 9 |
| Ankle angle | -0.98 | -0.28 to -1.69 | 74% | 7 | 5 |
| Nerve conduction velocity | -2.20 | -1.66 to -2.75 | 73% | 19 | 18 |
| Amplitude | -0.79 | 0.11 to -1.69 | 83% | 9 | 9 |
| Axon count | -1.40 | -0.78 to -2.01 | 86% | 27 | 18 |
| Diameter | -1.10 | -0.32 to -1.89 | 82% | 13 | 11 |

### Subgroup analyses

Subgroup analysis of all subgroups containing a minimum of 5 comparisons revealed a significant difference in muscle weight when comparing graft length between acellular allografts and conventional nerve autografts. Autografts showed a more favorable result in long grafts (> 2 cm) than in medium and short grafts (0-1 cm, > 1-2 cm) compared to acellular allografts (see Table 3).

**Table 3.** Subgroup analysis for graft length.

| Outcome measurement | SMD (Hedges g) | 95% confidence interval | P-value | No. of comparisons |
| --- | --- | --- | --- | --- |
| Muscle weight |  |  |  |  |
| - Short vs. long | -2.13 vs. -4.21 | -1.33 to -2.94 vs. -2.87 to -5.55 | 0.045 | 18 vs. 9 |
| - Medium vs. long | -1.92 vs. -4.21 | -1.09 to -2.75 vs. -2.87 to -5.55 | 0.026 | 18 vs. 9 |
| Nerve conduction velocity |  |  |  |  |
| - Short vs. long | -2.36 vs. -1.92 | -1.49 to -3.23 vs. -0.88 to -2.95 | 1 | 8 vs. 6 |
| - Medium vs. long | -2.33 vs. -1.92 | -1.23 to -3.43 vs. -0.88 to -2.95 | 1 | 5 vs. 6 |
| Axon count |  |  |  |  |
| Short vs. long | -1.46 vs. -1.21 | -0.54 to -2.38 vs. -0.08 to -2.33 | 1 | 13 vs. 9 |
| Medium vs. long | -1.62 vs. -1.21 | -0.14 to -3.11 vs. -0.08 to -2.33 | 1 | 5 vs. 9 |

However, for nerve conduction velocity and axon count no significant difference was found comparing graft length between acellular allografts and conventional nerve autografts.

All other subgroup analyses on graft length could not be interpreted because groups consisted of fewer than 5 studies. The same goes for all subgroup analyses for species.

### Sensitivity analysis

Exclusion of the studies published before 2008 did not alter our results significantly (see S2 Table). Also, when the studies were excluded in which animals were their own control no significant changes were found, only the amplitude SMD improved significantly (0.70 to 1.00), in favor of autografts (S3 Table).

Conclusions of all subgroup analyses appeared to be robust.

### Publication bias analysis

Publication bias could only be assessed for axonal count, muscle weight and nerve conduction velocity, because all other outcome measurements consisted of fewer than 15 independent studies. The funnel plot for muscle weight and axon count suggested some asymmetry. Duval and Tweedie’s Trim and Fill analysis resulted in 14 and 6 extra data points (Fig 5,6), indicating the presence of publication bias and some overestimation of the identified summary effect size. No publication bias for nerve conduction velocity was found (Fig 7).

**Fig 5.**
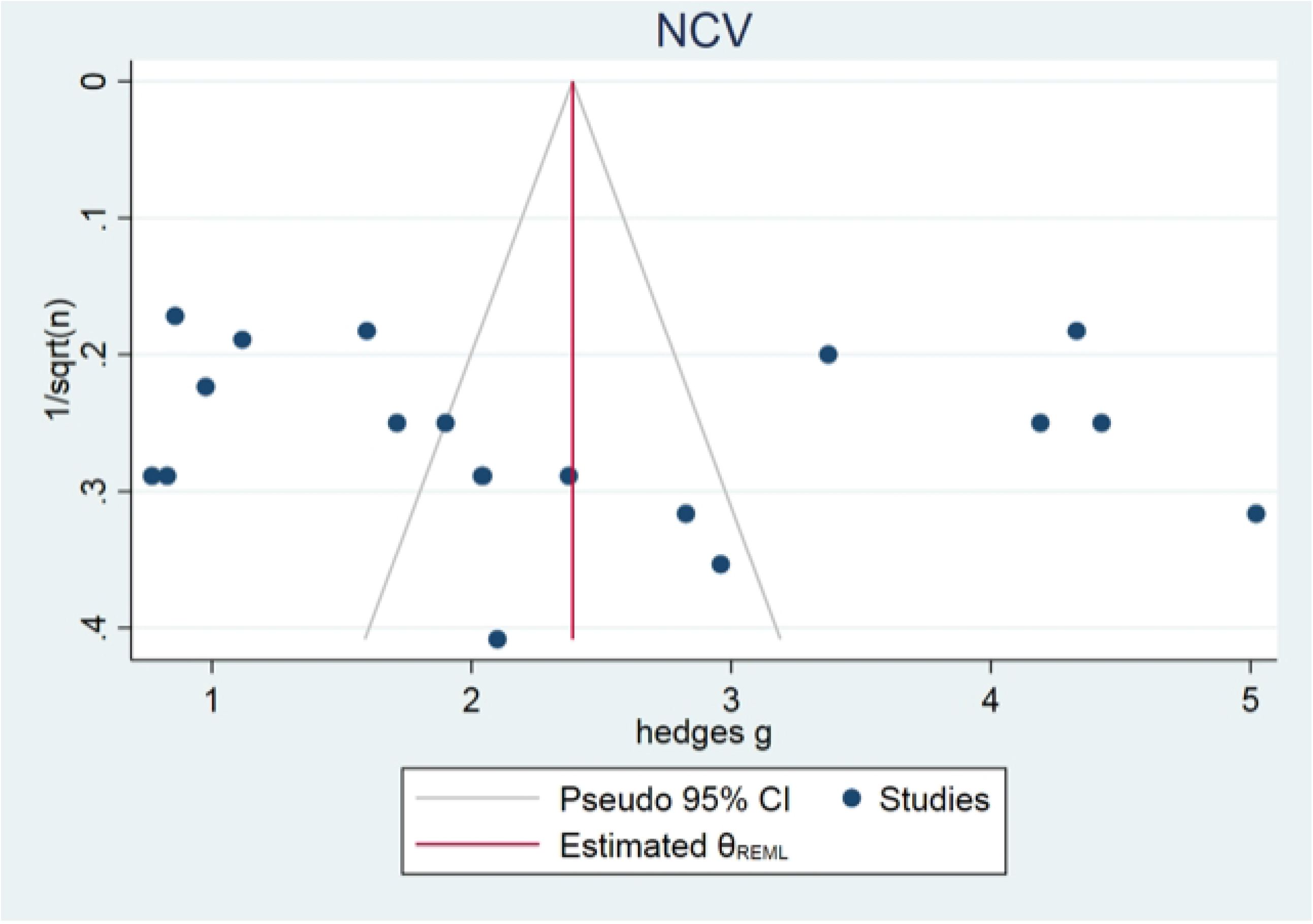
Muscle weight publication bias.

**Fig 6.**
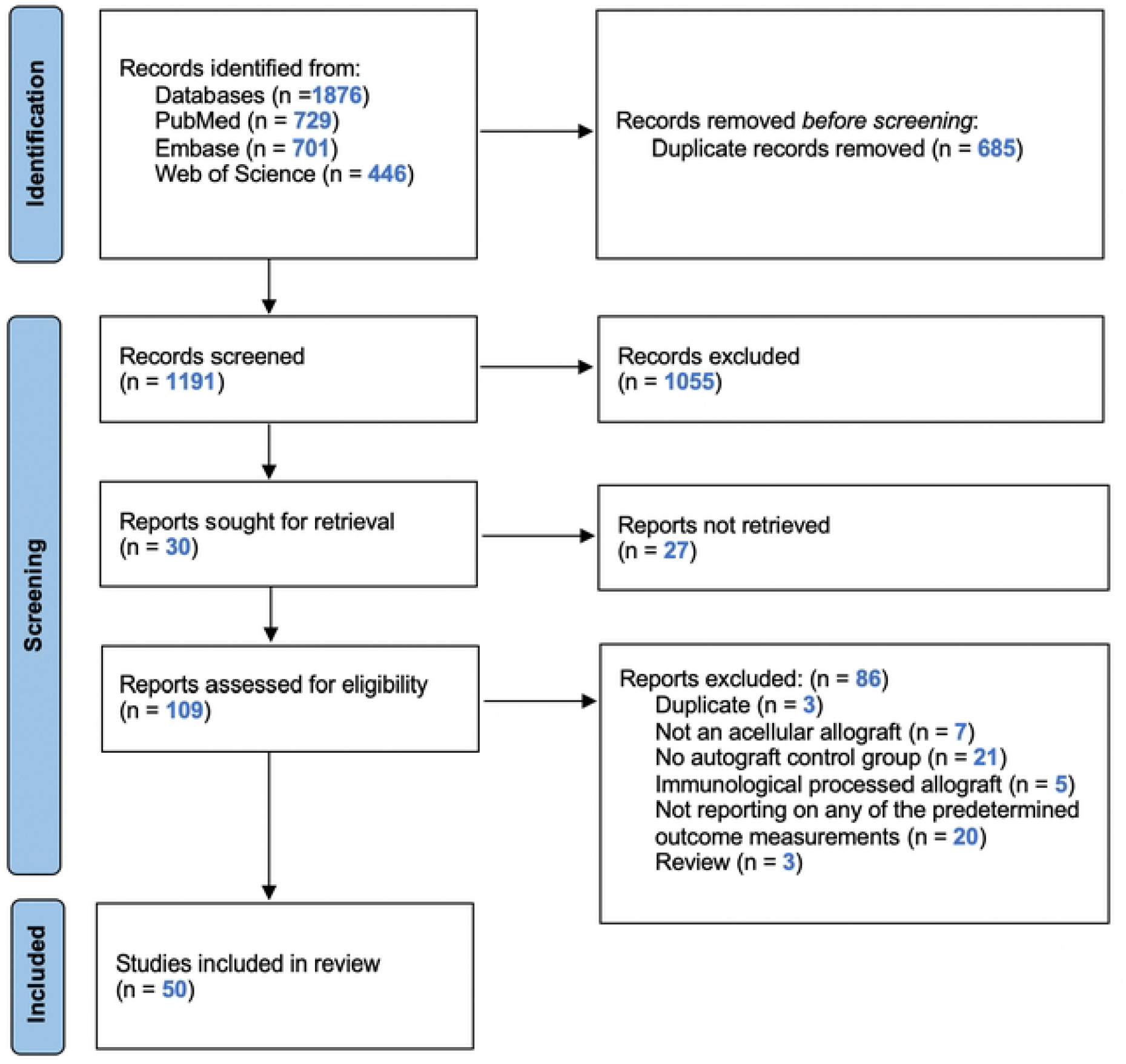
Axon count publication bias.

**Fig 7.**
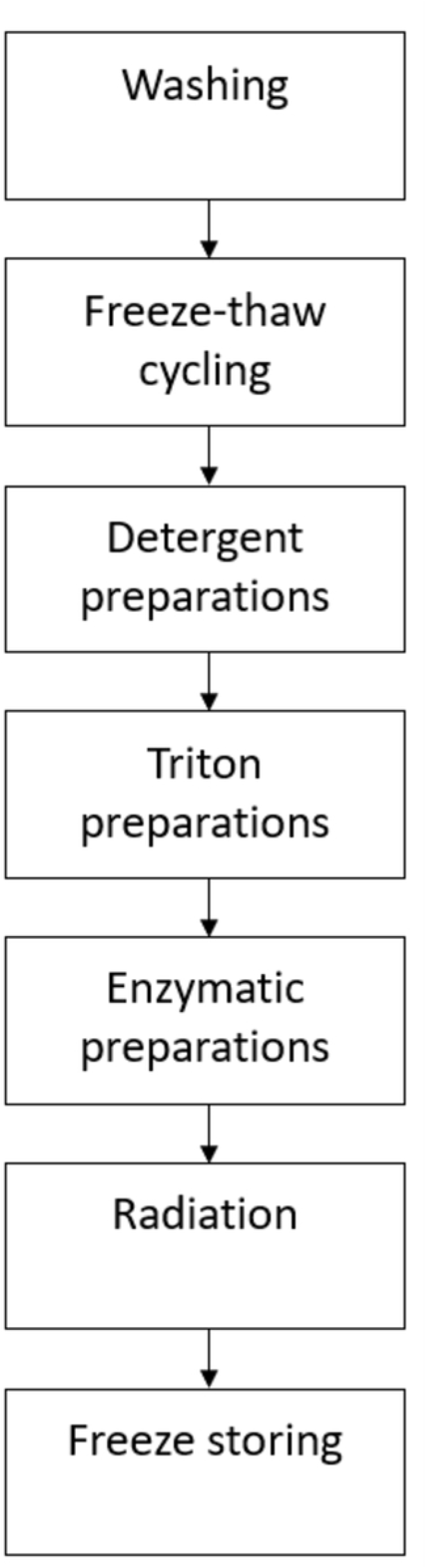
Nerve conduction velocity publication bias.

## Discussion

This review demonstrates that an acellular nerve allograft results in a significantly inferior nerve recovery compared to an autograft in animal models. Our subgroup analysis suggests that when an allograft is being used an allograft in short and medium (0-1cm, > 1-2cm) nerve gaps is preferred over an allograft in long (> 2cm) nerve gaps.

Comparing the available literature regarding the use of acellular allografts was challenging because a large variation in decellularization techniques were used. A variety of methods have been studied to prepare an acellular allograft by these labs such as cold preservation, freeze-thaw cycling, chemical detergent and enzymatic preparations, lyophilization and irradiation. (15-17) There is only one such method that is FDA approved available to surgeons produced by AxoGen, Inc., Alachua, Florida. AxoGen develops its human allografts by combining proprietary detergent processing and gamma irradiation which removes cellular remnants while minimizing the microstructural damage. Next to that, chondroitin sulfate proteoglycan is enzymatic removed from the endoneural tube system to advance axon regeneration. (18) We noticed that the studies we included, published before 2008, used a minimal decellularization method like mere freeze-thaw cycling. Based on the available data, no clear statement could be made as to which decellularization method is superior.

A few human studies show a resemblance in effect between the acellular allograft and the autograft. This would be a significant development because the allograft has a couple of fundamental advantages as opposed to the autograft. It has an unlimited supply that offers an excellent solution, e.g., plexus surgery after a major trauma. In such trauma, there can be insufficient autograft material to repair the nerve deficits. Therefore, in these cases the allograft offers a solution to restore the damages done. It also avoids potential donor site morbidity such as pain and loss of sensation. Next to that, there is the benefit of a shorter operation time. And finally, the off-the shelf availability of the allograft.

The clinical use of commercially available human acellular nerve allografts (AxoGen) for nerve reconstruction has been reported in several case reports and in a more sizeable multicenter study (RANGER). (13, 72-76) Unfortunately, the multicenter study lacks the opportunity to say anything about the effectiveness of the allograft, because it did not compare it to an autograft.

Safa et al. (75) and Leckenby et al. (76) both reported data from the RANGER study. Safa et al. conducted an analysis of 365 patients with 624 nerve repairs (AxoGen). They found a meaningful sensible and motor function recovery in 82% of cases. Finally, the authors stated that nerve defects up to 7 cm could achieve a useful recovery after treatment. Leckenby et al. analyzed 171 nerve repairs (AxoGen) in 129 subjects exploring a meaningful sensory and motor function recovery as well. In 73.7% of their cases a meaningful sensory recovery (^3^S3) was achieved. The percentage of meaningful motor recovery (^3^M3) was lower at 40,1%, respectively.

Neubauer et al. (18) observed another interesting aspect. They investigated which type of acellular nerve, i.e. motor, sensory or mixed type, is best used for repairing particular types of nerve gaps. No significant differences were found when comparing these three decellularized nerve types as a dominant grafting with regard to axon count and myelinated axons. Most repaired nerves happened to be a sensory dominant one in this study. We noticed that in the animal experiment the opposite is used. The use of motor types was rather common.

Until this day there is no proper “gold standard” to test nerve recovery, although the ultimate goal of nerve recovery is to maximize sensation and motion. The most commonly used outcome measurement for sensation is the von Frey test. (77) For motion, walking track analysis was believed to be the best overall assessment. (23, 78, 79) It is rarely used and some would say it is even obsolete. Additionally, walking track analysis does not reflect maximum muscle force capacity. Others say the most precise measurement is the isometric response of muscle to tetanic contraction. (80) The authors are aware that histomorphometry, electrophysiology and axonal count in particular may have a limited correlation to the real functional recovery of sensation and strength. (81) Next to that, histomorphometry is difficult to compare between different laboratories, because other methods to measure the outcome were used. We used a standardized mean difference for our meta-analysis to compensate for these differences. Over the years methods have evolved from manually calculating axonal count from a light microscopic photograph to a computer calculated estimate. The methods used by the studies in this review vary as well. Searching the publication databases, we found little evidence on which one is the best or on a clear sensitivity or specificity for these methods. However, Kim et al. (82) concluded that the semi-automated method for counting axons in transmission electron microscopic images was strongly correlated with conventional counting methods and showed excellent reproducibility. Nevertheless, the techniques for histomorphometry will always be an estimation and therefore prone to bias.

### Limitations of this review

Firstly, the risk of bias analysis revealed that many essential methodological details were poorly reported in the majority of included studies, which is why most risk of bias items assessed in this analysis were scored as ‘unclear risk of bias’. Drawing reliable conclusions from the included animal studies may have hampered the reliability of the analyses presented in this review.

Secondly, for some outcome measurements the number of included studies in this meta-analysis is relatively low, as a consequence the results of these small meta analyses may be imprecise. Next to that, the heterogeneity between the studies was moderate to high. We used a random effects model, subgroup analyses and conducted two different sensitivity analyses to account for this anticipated heterogeneity.

## Conclusion

Our review shows that the several different animal experiments, investigating the use of acellular allografts, are difficult to compare due to the wide variety of study designs used and the generally poor reporting of essential methodological details. Our results indicate that treating a nerve gap with an allograft results in an inferior nerve recovery compared to an autograft in seven out of eight outcome measurements assessed in animal models. In addition, we suggest that when an allograft is being used an allograft in short and medium (0-1cm, > 1-2cm) nerve gaps performs better than an allograft in long (> 2cm) nerve gaps. Large population studies or comparative clinical trials with large study populations to allow for participant and injury heterogeneity will be needed to prove and improve the success of the acellular allograft. We strongly advise future animal studies to be designed and reported according to the ARRIVE guidelines. (85, 86).

## Acknowledgements

The authors would like to thank Mrs On Ying Chan (Health Sciences reference librarian, Radboud University) for assisting with the development of the search strategy.

## Supporting information

**S1 Table. Search strategy**.

**S2 Table. Sensitivity analyses for exclusion of the studies published before 2008**.

**S3 Table. Sensitivity analysis for exclusion of studies in which animals were their own control**.

**S1 Fig. Raw Data**.

## Notes

### Competing Interest Statement

The authors have declared no competing interest.

## References

1. Sunderland S. Nerve and Nerve Injuries. Edinburgh: Livingstone. 1978.

2. Kline DGH, A. R Nerve Injuries: Operative Results for Major Nerve Injuries, Entrapments and Tumors Philadelphia: WB Saunders. 1995.

3. Noble J, Munro CA, Prasad VS, Midha R. Analysis of upper and lower extremity peripheral nerve injuries in a population of patients with multiple injuries. J Trauma. 1998;45(1):116–22.

4. Robinson PP, Loescher AR, Smith KG. A prospective, quantitative study on the clinical outcome of lingual nerve repair. Br J Oral Maxillofac Surg. 2000;38(4):255–63.

5. Jarvis MF, Boyce-Rustay JM. Neuropathic pain: models and mechanisms. Curr Pharm Des. 2009;15(15):1711–6.

6. Berger A, Millesi H. Nerve grafting. Clin Orthop Relat Res. 1978(133):49–55.

7. Stang F, Stollwerck P, Prommersberger KJ, van Schoonhoven J. Posterior interosseus nerve vs. tmedial cutaneous nerve of the forearm: differences in digital nerve reconstruction. Arch Orthop Trauma Surg. 2013;133(6):875–80.

8. Ehretsman RL, Novak CB, Mackinnon SE. Subjective recovery of nerve graft donor site. Ann Plast Surg. 1999;43(6):606–12.

9. Bamba R, Loewenstein SN, Adkinson JM. Donor site morbidity after sural nerve grafting: A systematic review. J Plast Reconstr Aesthet Surg. 2021;74(11):3055–60.

10. Daly W, Yao L, Zeugolis D, Windebank A, Pandit A. A biomaterials approach to peripheral nerve regeneration: bridging the peripheral nerve gap and enhancing functional recovery. J R Soc Interface. 2012;9(67):202–21.

11. Shin RH, Friedrich PF, Crum BA, Bishop AT, Shin AY. Treatment of a segmental nerve defect in the rat with use of bioabsorbable synthetic nerve conduits: a comparison of commercially available conduits. J Bone Joint Surg Am. 2009;91(9):2194–204.

12. Dellon AL, Mackinnon SE. An alternative to the classical nerve graft for the management of the short nerve gap. Plast Reconstr Surg. 1988;82(5):849–56.

13. Brooks DN, Weber RV, Chao JD, Rinker BD, Zoldos J, Robichaux MR, et al. Processed nerve allografts for peripheral nerve reconstruction: a multicenter study of utilization and outcomes in sensory, mixed, and motor nerve reconstructions. Microsurgery. 2012;32(1):1–14.

14. Banegas R. Nerve allografts - a review. BMC Proceedings 9. 2015.

15. Hiles RW. Freeze dried irradiated nerve homograft: a preliminary report. Hand. 1972;4(1):79–84.

16. Myckatyn TM, Mackinnon SE. A review of research endeavors to optimize peripheral nerve reconstruction. Neurol Res. 2004;26(2):124–38.

17. Sondell M, Lundborg G, Kanje M. Regeneration of the rat sciatic nerve into allografts made acellular through chemical extraction. Brain Res. 1998;795(1-2):44–54.

18. Neubauer D, Graham JB, Muir D. Chondroitinase treatment increases the effective length of acellular nerve grafts. Exp Neurol. 2007;207(1):163–70.

19. Hooijmans CR, Tillema A, Leenaars M, Ritskes-Hoitinga M. Enhancing search efficiency by means of a search filter for finding all studies on animal experimentation in PubMed. Lab Anim. 2010;44(3):170–5.

20. de Vries RB, Hooijmans CR, Tillema A, Leenaars M, Ritskes-Hoitinga M. A search filter for increasing the retrieval of animal studies in Embase. Lab Anim. 2011;45(4):268–70.

21. Ouzzani M, Hammady H, Fedorowicz Z, Elmagarmid A. Rayyan-a web and mobile app for systematic reviews. Syst Rev. 2016;5(1):210.

22. Hooijmans CR, Rovers MM, de Vries RB, Leenaars M, Ritskes-Hoitinga M, Langendam MW. SYRCLE’s risk of bias tool for animal studies. BMC Med Res Methodol. 2014;14:43.

23. Hadlock TA, Koka R, Vacanti JP, Cheney ML. A comparison of assessments of functional recovery in the rat. J Peripher Nerv Syst. 1999;4(3-4):258–64.

24. Boriani F, Savarino L, Fazio N, Pedrini FA, Fini M, Nicoli Aldini N, et al. Auto-Allo Graft Parallel Juxtaposition for Improved Neuroregeneration in Peripheral Nerve Reconstruction Based on Acellular Nerve Allografts. Ann Plast Surg. 2019;83(3):318–25.

25. Cai M, Huang T, Hou B, Guo Y. Role of Demyelination Efficiency within Acellular Nerve Scaffolds during Nerve Regeneration across Peripheral Defects. Biomed Res Int. 2017;2017:4606387.

26. Chato-Astrain J, Philips C, Campos F, Durand-Herrera D, Garcia-Garcia OD, Roosens A, et al. Detergent-based decellularized peripheral nerve allografts: An in vivo preclinical study in the rat sciatic nerve injury model. Journal of Tissue Engineering and Regenerative Medicine. 2020;14(6):789–806.

27. Agudo M, Woodhoo A, Webber D, Mirsky R, Jessen KR, McMahon SB. Schwann cell precursors transplanted into the injured spinal cord multiply, integrate and are permissive for axon growth. Glia. 2008;56(12):1263–70.

28. Frerichs O, Fansa H, Schicht C, Wolf G, Schneider W, Keilhoff G. Reconstruction of peripheral nerves using acellular nerve grafts with implanted cultured Schwann cells. Microsurgery. 2002;22(7):311–5.

29. Gao ST, Zheng Y, Cai QQ, Deng ZS, Yao WT, Wang JQ, et al. Combination of Acellular Nerve Graft and Schwann Cells-Like Cells for Rat Sciatic Nerve Regeneration. Neural Plasticity. 2014;2014.

30. Giusti G, Lee JY, Kremer T, Friedrich P, Bishop AT, Shin AY. The influence of vascularization of transplanted processed allograft nerve on return of motor function in rats. Microsurgery. 2016;36(2):134–43.

31. Giusti G, Willems WF, Kremer T, Friedrich PF, Bishop AT, Shin AY. Return of motor function after segmental nerve loss in a rat model: comparison of autogenous nerve graft, collagen conduit, and processed allograft (AxoGen). J Bone Joint Surg Am. 2012;94(5):410–7.

32. Gulati AK, Cole GP. Nerve graft immunogenicity as a factor determining axonal regeneration in the rat. J Neurosurg. 1990;72(1):114–22.

33. Haase SC, Rovak JM, Dennis RG, Kuzon Jr WM, Cederna PS. Recovery of muscle contractile function following nerve gap repair with chemically acellularized peripheral nerve grafts. Journal of Reconstructive Microsurgery. 2003;19(4):241–8.

34. Huang HT, Xiao HX, Liu HW, Niu Y, Yan RZ, Hu M. A comparative study of acellular nerve xenografts and allografts in repairing rat facial nerve defects. Molecular Medicine Reports. 2015;12(4):6330–6.

35. Hu M, Zhang L, Niu Y, Xiao H, Tang P, Wang Y. Repair of whole rabbit facial nerve defects using facial nerve allografts. J Oral Maxillofac Surg. 2010;68(9):2196–206.

36. Hu J, Zhu QT, Liu XL, Xu YB, Zhu JK. Repair of extended peripheral nerve lesions in rhesus monkeys using acellular allogenic nerve grafts implanted with autologous mesenchymal stem cells. Exp Neurol. 2007;204(2):658–66.

37. Hundepool CA, Bulstra LF, Kotsougiani D, Friedrich PF, Hovius SER, Bishop AT, et al. Comparable functional motor outcomes after repair of peripheral nerve injury with an elastase-processed allograft in a rat sciatic nerve model. Microsurgery. 2018;38(7):772–9.

38. Ide C, Tohyama K, Tajima K, Endoh K, Sano K, Tamura M, et al. Long acellular nerve transplants for allogeneic grafting and the effects of basic fibroblast growth factor on the growth of regenerating axons in dogs: a preliminary report. Exp Neurol. 1998;154(1):99–112.

39. Jiang L, Zheng Y, Chen O, Chu T, Ding J, Yu Q. Nerve defect repair by differentiated adipose-derived stem cells and chondroitinase ABC-treated acellular nerves. Int J Neurosci. 2016;126(6):568–76.

40. Li Z, Peng J, Wang G, Yang Q, Yu H, Guo Q, et al. Effects of local release of hepatocyte growth factor on peripheral nerve regeneration in acellular nerve grafts. Exp Neurol. 2008;214(1):47–54.

41. Li XFZ, J. M.; Qin, Y. W.; Luo, S. X.; Cheng, J. W.;. Comparison of acellular nerve grafts prepared with three different methods. Chinese Journal of Tissue Engineering Research 2013;16(47):8817–20.

42. Moore AM, MacEwan M, Santosa KB, Chenard KE, Ray WZ, Hunter DA, et al. Acellular nerve allografts in peripheral nerve regeneration: a comparative study. Muscle Nerve. 2011;44(2):221–34.

43. Muheremu A, Wang XY, Huang ZY, Zhang F, Peng J, Ao Q. Combination of Acellular Nerve Allograft and Human Umbilical Wharton Jell Stem Cells to Bridge Rat Femoral Nerve Defect. Journal of Biomaterials and Tissue Engineering. 2016;6(1):79–84.

44. Nakamoto JC, Wataya EY, Nakamoto HA, Santos GB, Ribaric I, Herrera AKA, et al. Evaluation of the Use of Nerve Allograft Preserved in Glycerol. Plast Reconstr Surg Glob Open. 2021;9(4):e3514.

45. Piao C, Li Z, Ding J, Kong D. Mechanical properties of the sciatic nerve following combined transplantation of analytically extracted acellular allogeneic nerve and adipose-derived mesenchymal stem cells. Acta Cir Bras. 2020;35(4):e202000405.

46. Qiu S, Rao Z, He F, Wang T, Xu Y, Du Z, et al. Decellularized nerve matrix hydrogel and glial-derived neurotrophic factor modifications assisted nerve repair with decellularized nerve matrix scaffolds. J Tissue Eng Regen Med. 2020;14(7):931–43.

47. Rovak JM, Mungara AK, Aydin MA, Cederna PS. Effects of vascular endothelial growth factor on nerve regeneration in acellular nerve grafts. J Reconstr Microsurg. 2004;20(1):53–8.

48. Saheb-Al-Zamani M, Yan Y, Farber SJ, Hunter DA, Newton P, Wood MD, et al. Limited regeneration in long acellular nerve allografts is associated with increased Schwann cell senescence. Exp Neurol. 2013;247:165–77.

49. Shin YH, Park SY, Kim JK. Comparison of systematically combined detergent and nuclease-based decellularization methods for acellular nerve graft: An ex vivo characterization and in vivo evaluation. J Tissue Eng Regen Med. 2019;13(7):1241–52.

50. Sun XH, Che YQ, Tong XJ, Zhang LX, Feng Y, Xu AH, et al. Improving nerve regeneration of acellular nerve allografts seeded with SCs bridging the sciatic nerve defects of rat. Cell Mol Neurobiol. 2009;29(3):347–53.

51. Tang P, Kilic A, Konopka G, Regalbuto R, Akelina Y, Gardner T. Histologic and functional outcomes of nerve defects treated with acellular allograft versus cabled autograft in a rat model. Microsurgery. 2013;33(6):460–7.

52. Tang P, Whiteman DR, Voigt C, Miller MC, Kim H. No Difference in Outcomes Detected Between Decellular Nerve Allograft and Cable Autograft in Rat Sciatic Nerve Defects. J Bone Joint Surg Am. 2019;101(10):e42.

53. Vasudevan S, Huang J, Botterman B, Matloub HS, Keefer E, Cheng J. Detergent-free Decellularized Nerve Grafts for Long-gap Peripheral Nerve Reconstruction. Plast Reconstr Surg Glob Open. 2014;2(8):e201.

54. Wakimura Y, Wang W, Itoh S, Okazaki M, Takakuda K. An Experimental Study to Bridge a Nerve Gap with a Decellularized Allogeneic Nerve. Plast Reconstr Surg. 2015;136(3):319e–27e.

55. Wang W, Itoh S, Takakuda K. Comparative study of the efficacy of decellularization treatment of allogenic and xenogeneic nerves as nerve conduits. J Biomed Mater Res A. 2016;104(2):445–54.

56. Wang D, Liu XL, Zhu JK, Hu J, Jiang L, Zhang Y, et al. Repairing large radial nerve defects by acellular nerve allografts seeded with autologous bone marrow stromal cells in a monkey model. J Neurotrauma. 2010;27(10):1935–43.

57. Wang H, Wu J, Zhang X, Ding L, Zeng Q. Study of synergistic role of allogenic skin-derived precursor differentiated Schwann cells and heregulin-1β in nerve regeneration with an acellular nerve allograft. Neurochem Int. 2016;97:146–53.

58. Wang Y, Zhao Z, Ren Z, Zhao B, Zhang L, Chen J, et al. Recellularized nerve allografts with differentiated mesenchymal stem cells promote peripheral nerve regeneration. Neurosci Lett. 2012;514(1):96–101.

59. Whitlock EL, Tuffaha SH, Luciano JP, Yan Y, Hunter DA, Magill CK, et al. Processed allografts and type I collagen conduits for repair of peripheral nerve gaps. Muscle Nerve. 2009;39(6):787–99.

60. Xiang F, Wei D, Yang Y, Chi H, Yang K, Sun Y. Tissue-engineered nerve graft with tetramethylpyrazine for repair of sciatic nerve defects in rats. Neurosci Lett. 2017;638:114–20.

61. Yan Y, Wood MD, Hunter DA, Ee X, Mackinnon SE, Moore AM. The Effect of Short Nerve Grafts in Series on Axonal Regeneration Across Isografts or Acellular Nerve Allografts. J Hand Surg Am. 2016;41(6):e113–21.

62. Yu H, Peng J, Guo Q, Zhang L, Li Z, Zhao B, et al. Improvement of peripheral nerve regeneration in acellular nerve grafts with local release of nerve growth factor. Microsurgery. 2009;29(4):330–6.

63. Yu T, Wen L, He J, Xu Y, Li T, Wang W, et al. Fabrication and evaluation of an optimized acellular nerve allograft with multiple axial channels. Acta Biomater. 2020;115:235–49.

64. Zhang LX, Tong XJ, Sun XH, Tong L, Gao J, Jia H, et al. Experimental study of low dose ultrashortwave promoting nerve regeneration after acellular nerve allografts repairing the sciatic nerve gap of rats. Cell Mol Neurobiol. 2008;28(4):501–9.

65. Zhang YR, Zhang H, Katiella K, Huang WH. Chemically extracted acellular allogeneic nerve graft combined with ciliary neurotrophic factor promotes sciatic nerve repair. Neural Regeneration Research. 2014;9(14):1358–64.

66. Zhao Z, Wang Y, Peng J, Ren ZW, Zhang L, Guo QY, et al. Improvement in Nerve Regeneration Through a Decellularized Nerve Graft by Supplementation With Bone Marrow Stromal Cells in Fibrin. Cell Transplantation. 2014;23(1):97–110.

67. Zhao Z, Wang Y, Peng J, Ren ZW, Zhan SF, Liu Y, et al. REPAIR OF NERVE DEFECT WITH ACELLULAR NERVE GRAFT SUPPLEMENTED BY BONE MARROW STROMAL CELLS IN MICE. Microsurgery. 2011;31(5):388–94.

68. Zhong H, Chen B, Lu S, Zhao M, Guo Y, Hou S. Nerve regeneration and functional recovery after a sciatic nerve gap is repaired by an acellular nerve allograft made through chemical extraction in canines. J Reconstr Microsurg. 2007;23(8):479–87.

69. Zhou LN, Zhang JW, Liu XL, Zhou LH. Co-Graft of Bone Marrow Stromal Cells and Schwann Cells Into Acellular Nerve Scaffold for Sciatic Nerve Regeneration in Rats. J Oral Maxillofac Surg. 2015;73(8):1651–60.

70. Zhou X, He B, Zhu ZW, He XH, Zheng CB, Xu J, et al. ETIFOXINE PROVIDES BENEFITS IN NERVE REPAIR WITH ACELLULAR NERVE GRAFTS. Muscle & Nerve. 2014;50(2):235–43.

71. Zhu Z, Zhou X, He B, Dai T, Zheng C, Yang C, et al. Ginkgo biloba extract (EGb 761) promotes peripheral nerve regeneration and neovascularization after acellular nerve allografts in a rat model. Cell Mol Neurobiol. 2015;35(2):273–82.

72. Gunn S, Cosetti M, Roland JT, Jr. Processed allograft: novel use in facial nerve repair after resection of a rare racial nerve paraganglioma. Laryngoscope. 2010;120 Suppl 4:S206.

73. Guo Y, Chen G, Tian G, Tapia C. Sensory recovery following decellularized nerve allograft transplantation for digital nerve repair. J Plast Surg Hand Surg. 2013;47(6):451–3.

74. Dunn JC, Tadlock J, Klahs KJ, Narimissaei D, McKay P, Nesti LJ. Nerve Reconstruction Using Processed Nerve Allograft in the U.S. Military. Mil Med. 2021;186(5-6):e543–e8.

75. Safa B, Jain S, Desai MJ, Greenberg JA, Niacaris TR, Nydick JA, et al. Peripheral nerve repair throughout the body with processed nerve allografts: Results from a large multicenter study. Microsurgery. 2020;40(5):527–37.

76. Leckenby JI, Furrer C, Haug L, Juon Personeni B, Vögelin E. A Retrospective Case Series Reporting the Outcomes of Avance Nerve Allografts in the Treatment of Peripheral Nerve Injuries. Plast Reconstr Surg. 2020;145(2):368e–81e.

77. Kemp SW, Cederna PS, Midha R. Comparative outcome measures in peripheral regeneration studies. Exp Neurol. 2017;287(Pt 3):348–57.

78. Dellon AL, Mackinnon SE. Selection of the appropriate parameter to measure neural regeneration. Ann Plast Surg. 1989;23(3):197–202.

79. Koka R, Hadlock TA. Quantification of functional recovery following rat sciatic nerve transection. Exp Neurol. 2001;168(1):192–5.

80. Frykman GK, McMillan PJ, Yegge S. A review of experimental methods measuring peripheral nerve regeneration in animals. Orthop Clin North Am. 1988;19(1):209–19.

81. Wilbourn AJ. The electrodiagnostic examination with peripheral nerve injuries. Clin Plast Surg. 2003;30(2):139–54.

82. Kim CY, Rho S, Lee N, Lee CK, Sung Y. Semi-automated counting method of axons in transmission electron microscopic images. Vet Ophthalmol. 2016;19(1):29–37.

83. Macleod MR, van der Worp HB, Sena ES, Howells DW, Dirnagl U, Donnan GA. Evidence for the efficacy of NXY-059 in experimental focal cerebral ischaemia is confounded by study quality. Stroke. 2008;39(10):2824–9.

84. Hirst JA, Howick J, Aronson JK, Roberts N, Perera R, Koshiaris C, et al. The need for randomization in animal trials: an overview of systematic reviews. PLoS One. 2014;9(6):e98856.

85. Hooijmans CR, Leenaars M, Ritskes-Hoitinga M. A gold standard publication checklist to improve the quality of animal studies, to fully integrate the Three Rs, and to make systematic reviews more feasible. Altern Lab Anim. 2010;38(2):167–82.

86. Percie du Sert N, Hurst V, Ahluwalia A, Alam S, Avey MT, Baker M, et al. The ARRIVE guidelines 2.0: Updated guidelines for reporting animal research. PLoS Biol. 2020;18(7):e3000410.

